# Genome-wide expression profiling *Drosophila melanogaster* deficiency heterozygotes reveals diverse genomic responses

**DOI:** 10.1101/032920

**Authors:** Hangnoh Lee, Dong-Yeon Cho, Cale Whitworth, Robert Eisman, Melissa Phelps, John Roote, Thomas Kaufman, Kevin Cook, Steven Russell, Teresa M. Przytycka, Brian Oliver

## Abstract

Deletions, commonly referred to as deficiencies by Drosophila geneticists, are valuable tools for mapping genes and for genetic pathway discovery via dose-dependent suppressor and enhancer screens. More recently, it has become clear that deviations from normal gene dosage are associated with multiple disorders in a range of species including humans. While we are beginning to understand some of the transcriptional effects brought about by gene dosage changes and the chromosome rearrangement breakpoints associated with them, much of this work relies on isolated examples. We have systematically examined deficiencies on the left arm of chromosome *2* and characterize gene-by-gene dosage responses that vary from collapsed expression through modest partial dosage compensation to full or even over compensation. We found negligible long-range effects of creating novel chromosome domains at deletion breakpoints, suggesting that cases of changes in gene regulation due to altered nuclear architecture are rare. These rare cases include *trans* de-repression when deficiencies delete chromatin characterized as repressive in other studies. Generally, effects of breakpoints on expression are promoter proximal (~100 bp) or within the gene body. Genome-wide effects of deficiencies are observed at genes with regulatory relationships to genes within the deleted segments, highlighting the subtle expression network defects in these sensitized genetic backgrounds.

**Author summary:** Deletions alter gene dose in heterozygotes and bring distant regions of the genome into juxtaposition. We find that the transcriptional dose response is generally varied, gene-specific, and coherently propagates into gene expression regulatory networks. Analysis of deletion heterozygote expression profiles indicates that distinct genetic pathways are weakened in adult flies bearing different deletions even though they show minimal or no overt phenotypes. While there are exceptions, breakpoints have a minimal effect on the expression of flanking genes, despite the fact that different regions of the genome are brought into contact and that important elements such as insulators are deleted. These data suggest that there is little effect of nuclear architecture and long-range enhancer and/or silencer promoter contact on gene expression in the compact *Drosophila* genome.

## Introduction

*Deficiency (Df)* is a genetic definition for mutations that affect contiguous loci on a chromosome [1]. They are now known to be a result of DNA deletion [2] and have many important uses in genetic analysis. *Dfs* are part of an important series of tests for defining the nature of mutant alleles according to Muller's morphs [3] where, for example, an allele is said to be an amorph when, in the homozygous condition, it exhibits the same phenotype as when uncovered by a *Df* encompassing the locus. Genetic mapping by complementation tests using a series of defined *Dfs* is also common, although not necessarily definitive, since dose-dependent interactions between loci (non-allelic non-complementation) can also result in mutant phenotypes [2]. Many dominant dose-dependent suppressor and enhancer mutations had been identified in *Drosophila* by the 1930's [4] and screens for non-allelic modifiers of mutant phenotypes is one of the most important uses for large collections of *Dfs* that tile the genome. The genetic interactions uncovered in such screens can be extremely informative, since gene pairs showing dose-dependent interactions often encode near neighbors in genetic pathways or subunits of the same protein complex. *“Df* kit” screens for modifiers of a gene of interest can thus rapidly identify regions where genes encoding members of the same pathway reside [5]. However, despite the undisputed utility of *Dfs*, we know relatively little about how these widely used tools globally impact the transcriptome.

*Drosophila melanogaster* shows very little clear haploinsufficiency [2], with most mutant alleles recessive to the wild type allele. The largest group of haploinsufficient loci is the *Minutes*, which encode ribosomal proteins or elongation factors [6], suggesting that there is a very strong requirement for diploidy when it comes to ribosome biogenesis. However, like many other animals, *Drosophila* is sensitive to large-scale reductions in gene dose. In a classic study, the entire genome was examined for dosage effects using synthetic deletions generated through crosses between translocation-bearing flies [7] and this segmental aneuploidy screen demonstrated that, outside of haploinsufficent regions, deleterious effects of gene dose reduction are generally dependent on the amount of material removed rather than the particular locus. This pioneering work suggested that there are many small additive or cumulative effects of reduced gene dose and, as the extent of a deleted segment grows, more genes in any given pathway are perturbed [8]. Thus, the effects of dose alteration accumulate, propagate, and eventually collapse the network. The limit of approximately a 1% deletion of the genome that *Drosophila* tolerates [7] is likely to reflect the connectivity of the gene network and the limits of network robustness [8]. The small effects associated with dose reduction are the main reason that *Dfs* are so useful in enhancer and suppressor screens. The dose changes in pairs of genes close in a network result in a phenotype, even though dose reduction of either alone is without overt consequence.

With the more recent application of genomic approaches, we are beginning to understand more about the effect of gene dose on the expression of autosomal hemizygous (one-copy) genes in *Drosophila*. In general, gene expression goes down when gene dose is reduced, but not by 2-fold [8–16]: genes tend to be expressed at a higher level than expected if there were a simple one-to-one relationship between copy number and expression level. Such modest, but measurable, autosomal dosage compensation could be due to the biochemical properties of pathways and the regulatory interactions commonly found in molecular biology [17] or to a more global response to aneuploidy that specifically recognizes aneuploid segments and increases expression of all genes in that segment [12]. The latter is analogous to the sex chromosome dosage compensation system that globally increases expression of the single *X* in wild type *Drosophila* males [18]. There has been some debate as to whether the non-sex-chromosome (autosomal) dosage compensation response is due to a general effect, elevating the expression of all genes, or to a gene-by-gene effect consistent with classic gene regulation [8, 10]. Within the genome there are many genes that show a consistent response, which could be due to a general system, but there are also dramatic outliers, where expression of one-copy genes collapses or actually increases. The later are more consistent with disrupted positive or negative feedback loops. The best evidence for gene-by-gene regulatory compensation is the coherent propagation of expression changes across gene expression networks observed in *Df/+* flies [8]. The dose effects for essentially the entire genome have been probed in highly aneuploid *Drosophila* cell lines [15, 16], but the vast numbers of changes in these cell lines makes interpretation of propagation extremely challenging. Cell lines have also evolved copy number states and show variable degrees of dosage compensation, which confounds analysis. One way to help address issues relating to mechanisms of autosomal dosage compensation would be to obtain a larger sample of expression profiled *Df* genotypes. We have therefore examined the effects of chromosome arm *2L Dfs (Df(2L))* on transcription in adult females and males in two genetic backgrounds, generating a total of 815 expression profiles in biological duplicate (or greater).

We report on three aspects of the effect of *Df(2L)s* on the transcriptome. First, we show that one-copy gene expression is generally locus-specific, suggesting that biochemical processes and molecular regulatory circuits account for most autosomal dosage compensation. However, the genome is organized into chromatin domains flanked by insulators [19] and we also provide evidence that deletions within chromatin domains associated with repressive chromatin marks result in superior compensation or even overexpression. This counter-intuitive effect of increasing expression of one-copy genes suggests that there is a *trans* effect of *Dfs* that can weaken repressive domains. Surprisingly, these effects were preferentially found in females. Second, *Df* breakpoints bring together two regions of the genome that are usually distant in the linear chromosome. This can result in breakpoint proximal changes due to transcription unit fusions, or local changes due to juxtaposition of regulatory regions such as enhancers, and it may be expected that fusing domains *in cis* would generally result in altered expression within novel chromatin domains. In agreement with previous work on inversions [20], our results suggest that there is very little functional long-range promoter communication with enhancers or silencers, and that disrupting chromatin domains is generally innocuous in terms of transcription. Third, genes function in networks thus perturbations should act at distance in genomic or 3D nuclear space due to information propagation through dynamic biological systems controlled by transcriptional regulators. We find strong support for this type of network structure in the expression profiles, since we observed that reduction in transcript levels from one-copy genes propagates to primary network neighbors and is dissipated after tertiary network separation. This suggests that we can learn much about the logic of gene networks by measuring how they respond to dose changes in hemizygous conditions without overt phenotypes, rather than profiling mutants with morphological, physiological, or behavioral phenotypes that complicate pathway analysis.

## RESULTS

To systematically investigate the effects of deletions on transcription, we expression profiled a set of molecularly defined hemizygous DrosDel fly genotypes [21] uncovering approximately 68% of the euchromatic portion of the left arm of chromosome *2 (2L;* **Figure 1A**). We examined gene expression in adult females and males from 99 different *Df* genotypes following backcrosses to the *w^1118^* parental line used to generate the DrosDel collection. In addition, to determine how sensitive dosage responses were to genetic background, we also examined adult female and male gene expression in 67 of these *Df* genotypes in a hybrid background by outcrossing to the sequenced modENCODE *OregonR* line [15] for a total of 102 expression profiled *Dfs*. The hybrid background resulted in heterozygosity for ~636K single nucleotide polymorphisms (SNPs) and deletions or insertion (InDels) in addition to the heterozygosity associated with each *Df* (**Supplemental file 1**). Briefly, we performed multiplexed RNA-Seq on polyA+ selected mRNA, used ERCC spike-in controls [22] from two sub-pools to characterize measurement variance and ratiometric performance, and determined low expression cutoffs based on an evaluation of intergenic expression. At a minimum we used biological duplicates for each *Df* and each sex for a total of 815 expression profiles, which are available in the Gene Expression Omnibus (GEO, accession GSE61509 and GSE73920).

**Fig. 1.**
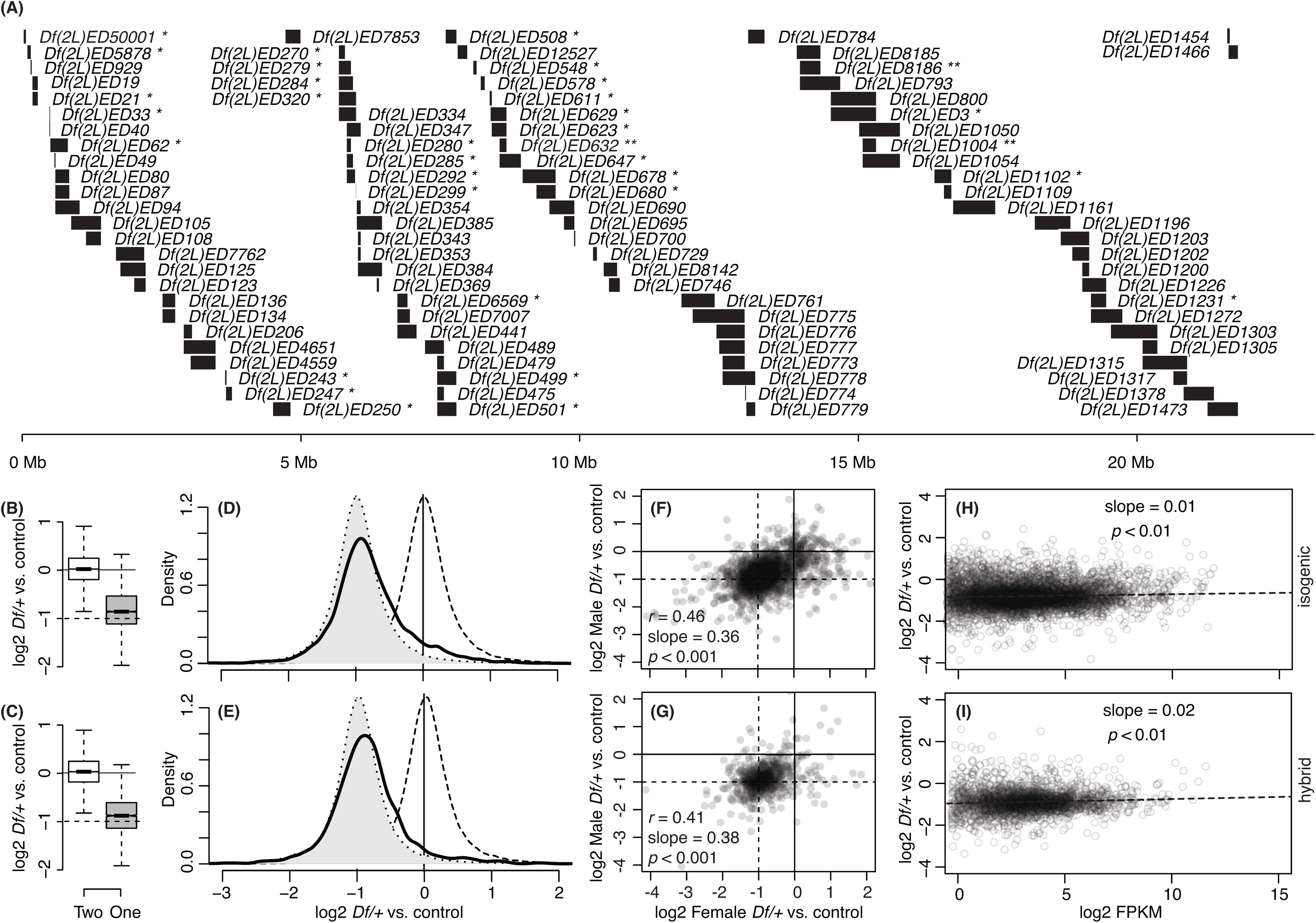
Transcriptional responses to gene-copy on Chromosome 2L. A) *Dfs* used in this study. The extent of the indicated deleted DNA (bars) and position along the first 25Mb of *2L* (bottom scale) are indicated. *Dfs* were tested in both the isogenic and hybrid background (except * isogenic only, and ** hybrid only). B, C) Boxplots of gene expression in two-copy genes (open) and one-copy genes (filled) relative to normalized global mean expression of the same genes in the rest of the dataset. Bottom, middle, and top lines of each box represent the 1^st^, 2^nd^ (median), and 3^rd^ quartile of the distribution. The maximum or minimum observation within 1.5 times of the interquartile range (3^rd^ quartile – 1^st^ quartile) from the 3^rd^ or 1^st^ quartiles is indicated by whiskers. Notches indicate the 95% confidence interval for the medians. D, E) Normalized relative expression value distributions of two-copy genes (dotted line, open), the projected distribution if gene expression were reduced by 50% in one-copy genes (dotted line, filled), and observed one-copy gene expression (solid line, open). F, G) Scatterplots that display one-copy gene expression levels between males and females from same *Dfs*. Pearson’s correlation coefficient (*r*) is indicated. H, I) One-copy versus two-copy gene expression levels plotted against the median expression levels of replicates expressed in units of <u>F</u>ragments <u>P</u>er <u>K</u>ilobase of transcript per <u>M</u>illion mapped reads (FPKM). B, D, F, H) Isogenic background. C, E, G, I) Hybrid background.

### Gene dose responses

To compare one-copy expression to two-copy expression for individual genes on *2L* we took advantage of the fact that there were many *Df/+* genotypes where a given gene was two-copy. Therefore, for any given gene, we took the median of two-copy gene expression in all lines as a reference for the expression when that gene was only in one-copy. To summarize the typical responses of genes to their own dose, we pooled the data for one-copy gene expression across all *Df/+* genotypes within the isogenic or hybrid backgrounds. In both genetic backgrounds, we observed a clear reduction in gene expression from one-copy *(p* < 0.001, Mann Whitney U test) as compared to two copies. However, our analysis confirmed previous reports that reduced expression is not 2-fold [8, 12–14]. We observed a mean 1.1-fold compensation in response to gene dose reduction (**Figure 1B,C**). As in previous work, we observed that compensation was not due to a uniform effect on all genes, as one-copy gene expression was skewed towards compensation (**Figure 1D,E**; Pearson’s second coefficient of skewness = 0.14–0.31 for one-copy genes compared to 0.01–0.03 for two-copy genes; kurtosis = 6.6–11.5 for one-copy genes compared to 12.8–14.4 for two-copy genes), with extended tails in the distributions of one-copy gene expression values (not shown in the truncated plots). These data indicate that different genes show differences in compensation responses. We observed similar (but not identical, as will be important later) compensation in females and males (**Figure 1F,G**), despite the highly dissimilar levels of expression between the sexes ([23], Materials and Methods). Thus, at least some of the response to copy number is a characteristic of an individual gene.

Increased compensation among genes showing low expression has been noted in several previous studies in *Drosophila melanogaster* [8, 13], which is unsurprising since genes with low expression are expected to be more sensitive to noise and therefore might require tighter expression level control. However, we observed no increased compensation for genes expressed at low levels in our study (**Figure 1H,I**) and this was independent of low-expression cut-off. Low gene expression in whole animals can be due to low uniform expression in most cells or high expression in limited cell types. Compensation has also been reported to be biased for broadly expressed genes [13]. We therefore asked if temporal or spatial heterogeneity, or representation of Gene Ontology (GO) terms correlated with compensation, but again we observed no significant trend (**not shown**). It is likely that data compression and increased contributions of technical noise to low-level gene expression measurements contributed to overestimating compensation at low expression levels in previous work and confounded subsequent analysis (see Discussion). Thus, while the dose response has a gene-specific component, we were not able to explain that response by particular gene expression levels or functional gene categories.

### Gene-specific dosage response examples

To further explore the influence of locus, sex and genetic background on the dosage response, we used overlapping *Dfs* to increase the number of expression measurements from one-copy genes. This analysis has the added advantage of determining if a particular *Df* used to uncover a gene altered the response. We examined a region near the middle of *2L* (cytological regions 33–34) with five distinct *Dfs*, and a second closer to the centromere (cytological regions 36–37) with four different *Dfs* (**Figure 2A-D**). We observed a complex variety of expression variance patterns, compensation responses, and sex- or allele-biased compensation depending on the individual locus. One-copy gene expression is generally noisier that two-copy gene expression, which we discuss at length elsewhere (Cho et al, companion paper), but this is also gene-specific. For example, we observed a wide ranges of responses to reducing the dose of *nubbin* (*nub*), from overcompensated (>2-fold increase) to anticompensation (>2-fold decrease), which also showed some genetic background-specificity, as we observed better compensation in the *modENCODE OregonR* background (**Figure 2A,B**). In contrast the *hook* gene showed no compensation across 24 different experiments (**Figure 2C,D**). We observed partial compensation of the *Multidrug-Resistance like Protein 1 (MRP)* locus in females (**Figure 2A**), but variable compensation in males (**Figure 2B**). We also observed a sex-biased response in the case of *Similar to deadpan* (*Sidpn*), which was overcompensated in females (**Figure 2C**) and partially compensated in males (**Figure 2D**). As expected, based on the correlation between compensation in females and males across *2L* (see **Figure 1F,G**), we found that many genes showed similar responses. For example, we observed over-compensation of *CG15485* (**Figure 2A,B**) and anti-compensation of *CG17572* (**Figure 2C,D**) in both sexes. Most ribosomal protein encoding genes are haploinsufficient, resulting in a Minute phenotype. We note with interest that the ribosomal-protein-encoding gene *RpL30* (**Figure 2C-E**) showed evidence of compensation, consistent with the lack of a Minute phenotype reported for mutations in this gene [6]. A second ribosomal protein encoding gene *(RpL7-like*, **Figure 2F**) also showed compensation. That these two loci are rare examples of ribosomal protein encoding genes that are not haplo-insufficient genetically and exceptionally well compensated at the transcriptional level supports the idea that stoichiometric mRNA levels of ribosomal protein encoding genes are ultimately important for ribosome function [6].

We observed one case where the particular uncovering *Df* correlated with a specific response. In males, the cluster of the *ACXA*, *ACXB*, *ACXC*, and *ACXE* genes showed very good compensation when uncovered by *Df(2L)ED775*, but much poorer compensation when uncovered by four other *Dfs* (**Figure 2AB**). The increased compensation when these genes were uncovered by *Df(2L)ED775* was also allele-specific as the effect was only observed in the isogenic background. The amount of DNA removed by *Df(2L)ED775* was more extensive than most of the *Dfs* used in the study (**Figure 1A**), raising the possibility that the extent of a deletion contributes to compensation. However, we observed no significant relationship between the length of hemizygous segments and dose responses in our experiments (**Figure 2G,H**).

**Fig 2.**
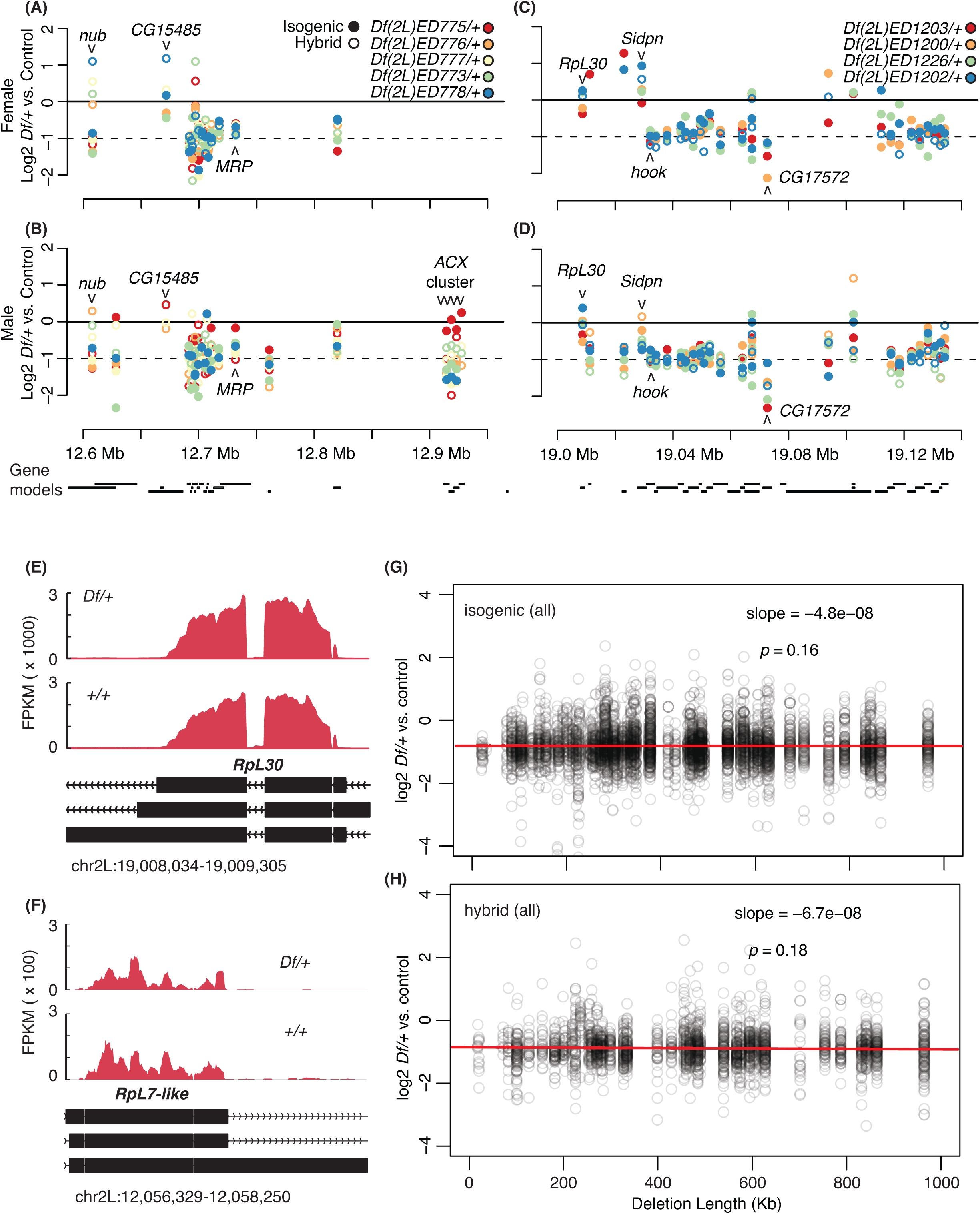
One-copy gene responses in different *Df/+* settings. A-D) One-copy versus two-copy gene expression levels plotted at the centers of each gene model (labeled below). Observations made in the isogenic genetic background (filled) and the hybrid genetic background (open) are shown. Genes whose expression is below the expression cutoff (see Methods) are not shown. A, B) One-copy gene expression of the genes between chr2L:12,545,800 and 12,975,028, which is uncovered by five different *Dfs*. C, D) One-copy gene expression of the genes between chr2L:19,003,398 and 19,158,447 region that is uncovered by four different *Dfs*. E, F) Sashimi plots that display normalized numbers of mapped reads across *RpL30* and *RpL7-like* gene body regions. Expression in *Df(2L)ED1202/+* for *RpL30* and *Df(2L)ED761/+* for *RpL7-like* was compared to expression in *Df(2L)ED774/+*, which was the shortest deletion in our study and is *+/+* for both genes. Exons (black bars) in the gene models and transcription direction (chevrons) are show below. G, H) Dosage responses (y-axis) of one-copy genes when uncovered by deletions of indicated length (x-axis). Results from the isogenic genetic background (top) and from the hybrid genetic background (bottom) are shown.

### Nuclear architecture and dosage responses

Our data do not support the idea that specific functional classes of genes or gene features, such as length, expression breadth or level, are associated with distinct dosage responses. However, we did notice that some blocks of genes showed common compensation responses and speculated that these might correspond to a particular chromatin state. For example, a group of genes [*CG18302, world cup (w-cup), CG31872, CG18284*, and *CG17097*, but not *Tripartite motif containing 9 (Trim9)*] uncovered by the proximal portion of *Df(2L)ED8142* showed overcompensation in females but not males, while the rest of the genes uncovered by this *Df* showed a more typical partial compensation response (**Figure 3A**). To explore the role of chromatin domains on the dosage response, we plotted our results along with the chromatin state maps from a DamID study on chromatin-associated protein occupancy ([24], **Figure 3B**), 3-D structure determination from Hi-C chromatin conformation capture mapping ([25], **Figure 3C**), and nuclear envelope attachment from a LaminB DamID study ([26], **Figure 3D**). In both the genetic backgrounds examined we observed dramatically improved dosage compensation in regions of the genome in structural domains associated with repressed gene expression (**Figure 3B,C**). These repressive domains show overlapping characteristics between the DamID and Hi-C studies, as well as being enriched in LaminB binding. When we specifically looked at LaminB domains, we also observed improved compensation (**Figure 3D**). We were very surprised to find that all these improved compensation distributions were only observed in females: we found no significant correlations, or even hints of a trend, between repressive chromatin and compensation in males. We also observed improved compensation in regions of Polycomb group (PcG) protein occupancy (**Figure 3B**), but not in the structural domains enriched in those proteins from Hi-C (**Figure 3C**). Again, we observed a correlation between PcG occupancy and compensation only in females. In addition to the increased median (and mean) compensation levels in these repressive chromatin domains, we observed an increased range of responses. Thus, there is a greater heterogeneity in the compensation response within these domains. We observed modest, but significant decreased compensation in one of the two types of active chromatin in both sexes based on occupancy (**Figure 3B**). Active regions of the “Yellow” type, which is enriched in H3K36me3 domains and in genes with broad expression patterns [24], showed significantly worse compensation, while active regions of the “Red” type showed the global dosage response. The locations of chromatin domains were defined from different samples (e.g. cells or embryos) than our RNA-Seq analysis (adult flies), but these data indicate that the female dosage response is different from males in contiguous regions regardless of underlying cause.

**Fig 3.**
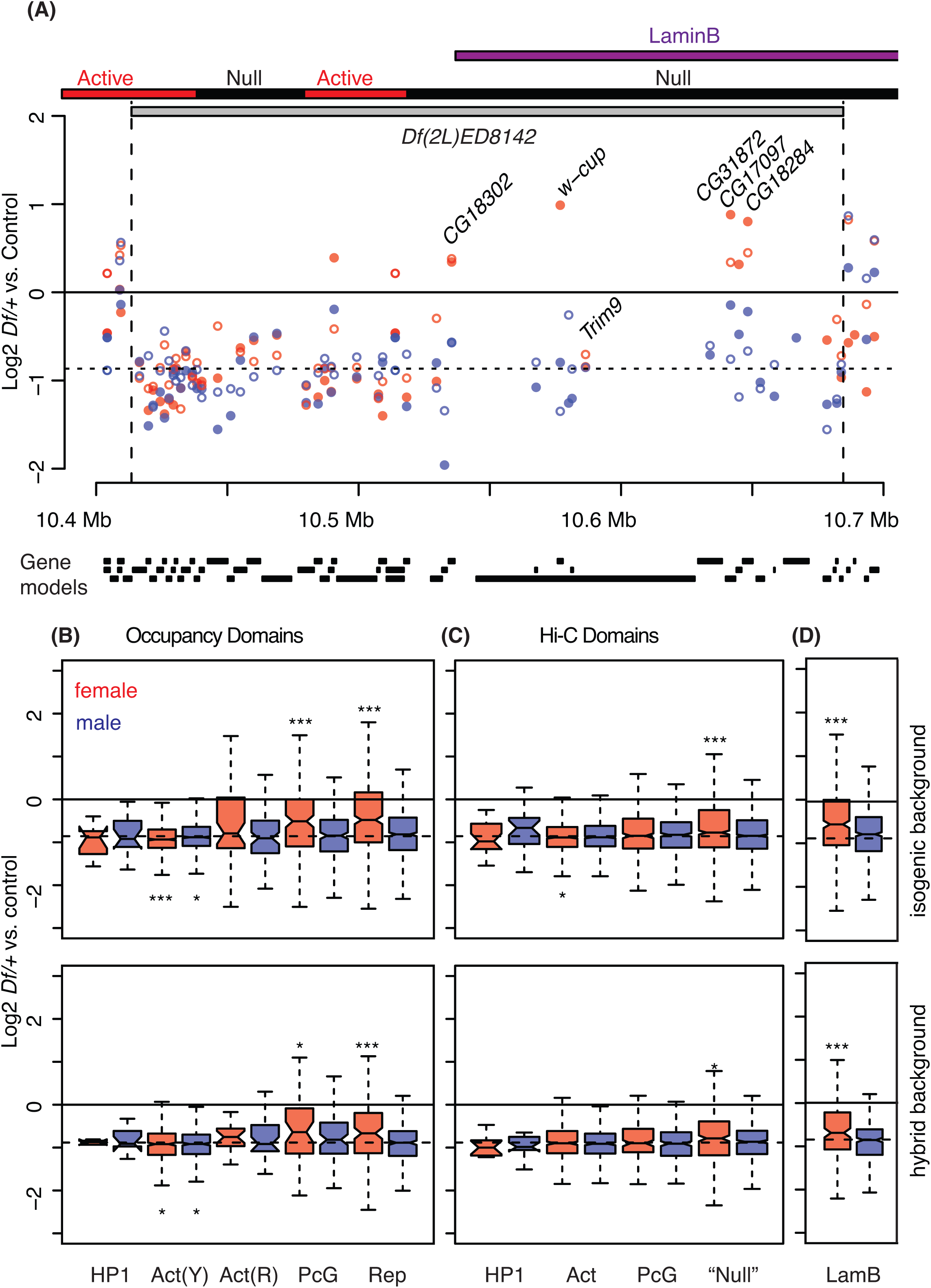
Disruption of chromatin and/or 3D nuclear domains by *Df* breakpoints. A) One-copy gene expression in *Df(2L)ED8142/+* plotted across the deletion position in log2 scale. Chromatin states from Hi-C [25] and LaminB domains from DamID [26] are presented above (bars). See Fig 2 for additional labeling. B-D) Autosomal dosage compensation levels were measured from one-copy genes that mapped to different chromatin state domains (B), topologically associated domains (C), and LaminB domains (D). Top panels. Data from the isogenic genetic background (top) and hybrid genetic background (bottom) are shown. Domains are labeled according to diagnostic enrichments/functions from the original studies: Heterochromatin Protein 1 domain (HP1), “yellow” [Act(Y)] and “red” [Act(R)] active domains, Polycomb Group domain (PcG), repressive domain (Rep), undefined and other (Null), and LaminB (LamB). * *p* < 0.05, ** *p* < 0.01, *** *p* < 0.001.

Due to the unexpected female-biased compensation of genes within repressive chromatin domains, we examined global sex-biased compensation to see if these blocks of improved compensation are revealed *in toto*. Indeed, while median and mean compensation levels were similar in females and males (**Figure 4A**), we observed more genes with very good compensation in females and more genes with intermediate compensation in males *(p* <0.05, Kolmogorov-Smirnov test; **Figure 4B**). When we asked if genes showed similar compensation between isogenic and hybrid backgrounds using expectation maximization clustering, we observed clear sets of genes with modest and full compensation in females, but we were unable to detect two clusters in males (**Figure 4C,D**). These data suggest that there are subtle differences in autosomal compensation between the sexes, as indicated by the correlation between repressive domains and female-biased compensation (Figure 3). Thus, at the level of a 2L-wide overview, our comparison between the sexes suggests varied gene-by-gene responses to copy reduction, consistent with a feedback/buffering model for dosage responses, with a contribution from chromatin structure in females. The increased compensation when repressive domains are deleted is consistent with a role for chromosome pairing and reinforcing repressive effects that are relaxed by deletion from one homolog.

**Fig 4.**
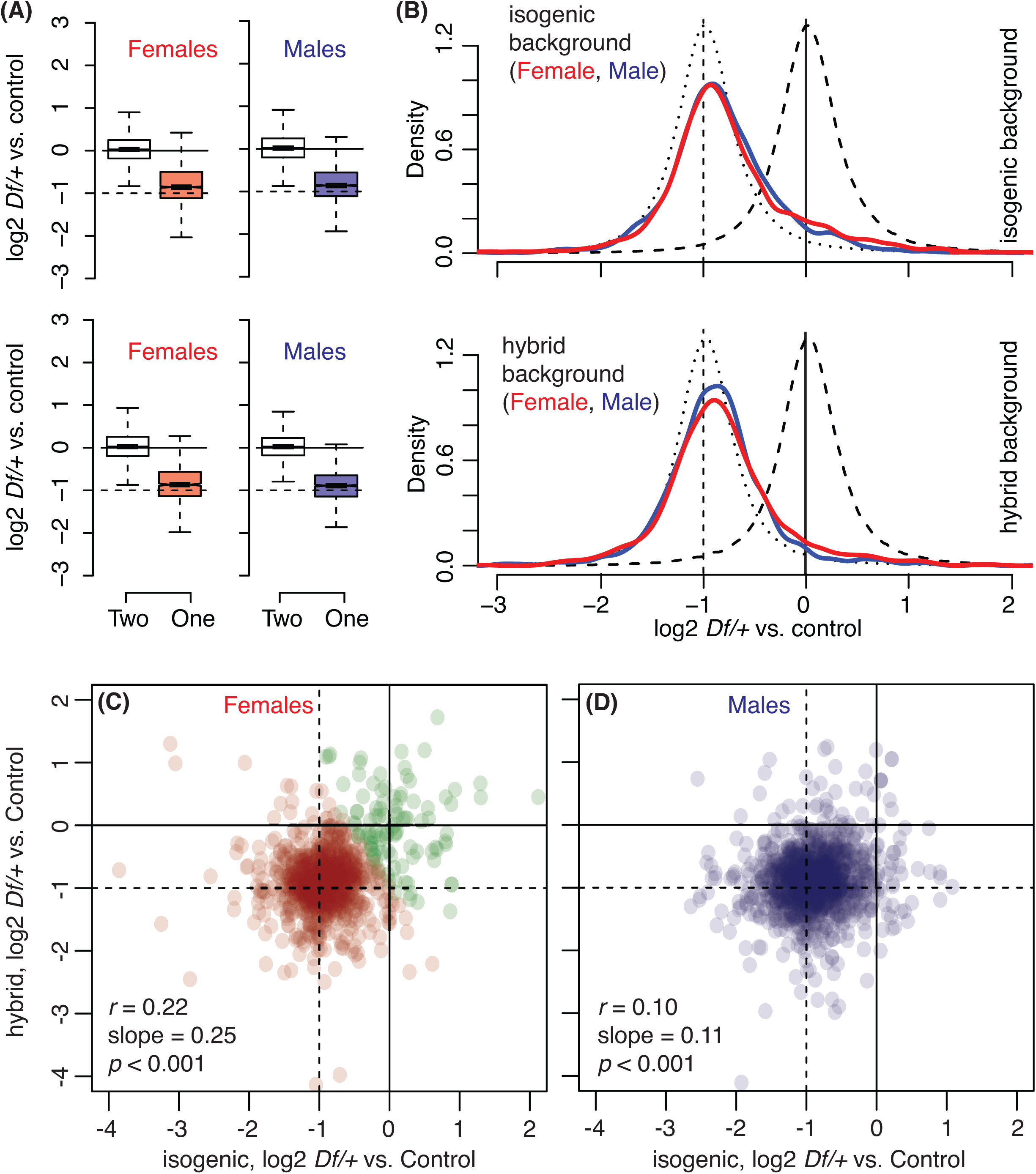
Sex-based difference in one-copy gene expression. A) Boxplots of gene expression in two-copy genes (light fill) or one-copy genes (red or blue fill) from all *Df* lines used (see Fig 1 for boxplot parameters). B) Density plots that display normalized relative expression value distributions of two-copy genes (dashed line), projected distribution if gene expression was reduced by 50% in one-copy genes (dotted line), and observed one-copy gene expression from females and males (solid red and blue). A, B) Data from isogenic (top) and hybrid (bottom) genetic backgrounds are shown. C, D) Scatterplots that compare one-copy gene expression relative to two-copy gene expression between the isogenic genetic background and hybrid genetic background in females (C) and males (D). A subset of genes in (C) represents “better compensated” genes identified by clustering analysis (Green). *r* = Pearson’s correlation coefficient. Slopes are from linear regression. P values are from F-tests.

### Breakpoints and gene expression

Deletions bring distant regions of the chromosome into linear juxtaposition: if such novel genome arrangements fuse domains or destroy insulator elements then they should create new expression environments. If repressive domains can be weakened by deletions, as suggested by our data in females, those effects might spread into adjacent regions of repressive chromatin that are juxtaposed with other types of chromatin. This is essentially the opposite of position effect variegation, where spreading of repressive chromatin is observed [27]. Our data was inconclusive as to whether active chromatin spreads into repressive domains, since we observed modest but significant effects on genes flanking breakpoints when the breakpoint was within a previously identified LamB repressive domain only in the hybrid background (**Figure 5A**). We also observed an ambiguous effect of breakpoints in HP1 domains on two copy gene expression, but only in females and the directionality of the change in expression differed in the two genetic backgrounds we assayed. We found slight and occasionally significant increased expression of two-copy genes flanking breakpoints in “Null” domains (**Figure 5A**). Breakpoints in other chromatin domains showed no significant changes in the expression of breakpoint proximal two copy genes. Our data suggests that LamB repressive domains are more sensitive to de-repression in *trans* than in *cis*. This suggests that LamB domain repression is additive or cooperative across homologs.

**Fig 5.**
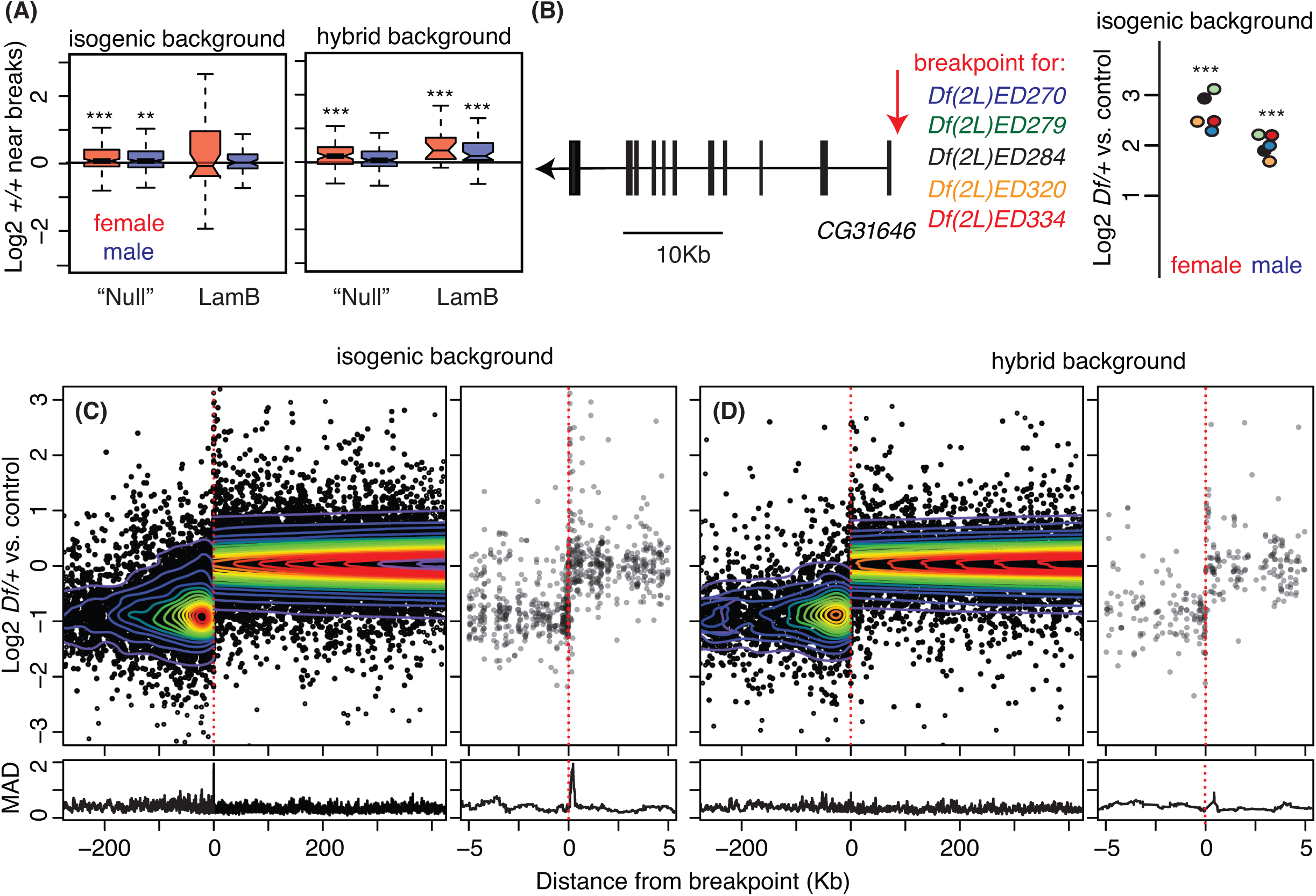
Disruption of DNA linear structure by *Df* breaks. A) Two-copy gene expression near *Df* breakpoints. The boxplots display distributions of normalized relative two-copy gene expression within the Null domain (from Hi-C), or LaminB domains that are disrupted by deletions (see Fig 1 for boxplot parameters and Fig 3 for statistics). B) A schematic of the *CG31646* gene model (left) showing the position of a common breakpoint (red arrow) for the 5 *Dfs*. Expression changes of CG31646 in different *Df* genotypes is also indicated (right). Colors of the filled circles match with the *Dfs* genotypes by the plot. *P* values (asterisk) are based on empirical Bayes moderated T-test in the limma package (See Methods, *** *p* < 0.001). C, D) Gene expression changes near breaks are collectively displayed by aligning all breakpoints in the study at “0” (red dotted line). One-copy genes are placed on the left side, and two-copy genes are place on the right side of 0. Contours represent data point density. Wide (left) and zoomed (right) views of the same data are shown. Variability in gene expression was summarized using Median Absolute Deviation (MAD, a non-parametric measure of the variability) from sliding windows of 30 genes (bottom).

Deletions can also remove *cis*-regulatory regions such as enhancers and silencers. For example *Df(2L)ED270, Df(2L)ED279, Df(2L)ED284, Df(2L)ED320*, and *Df(2L)ED334* all have a common breakpoint just upstream of the *CG31646* promoter, deleting a region where bearing a known CNS regulatory region [29] (**Figure 5B**). In males, and especially in females, these deletions resulted in dramatic overexpression of *CG31646* suggesting that a silencer was removed by each of these deletions. To determine if the effects of structural rearrangements on gene expression are common, we centered all the breakpoints from the *Dfs* used in this study and plotted expression flanking the breakpoint as well as the median absolute deviation to summarize the results. We observed no significant change in expression with distance from the breakpoint, with the exception of genes within 100bp of a breakpoint (**Figure 5C**). Even this breakpoint proximal effect is probably less significant than it appears, since the spike of increased expression in the isogenic background is due almost exclusively to the *Dfs* in **Figure 5B**. Thus despite the fact that the deletions our study removed 2,100 insulators identified in an embryo study [30], and have breakpoints that disrupt 437 chromatin domains identified in a Hi-C study [25], we find little evidence that these play major roles in transcription. These data suggest that hemizygous chromatin rarely alters expression on the deletion homolog and that bringing two separated regions into a novel configuration has little effect on the expression of two-copy genes near breakpoints. These data support the idea that the *Drosophila* genome is compact: most genes are regulated with promoter proximal regulatory sequences and show little long distance effects of chromatin structure. The regional effects of LamB domains in females are an exception.

### Propagation through gene networks

The general absence of breakpoint proximal effects of *Dfs* on transcription of two-copy genes does not mean that there is no effect of deletions on the rest of the genome. We observed tens of significant changes in gene expression for each gene made one-copy in a *Df/+* fly (**Supplemental file 2**, 3). However, the genes with changed expression were scattered across the genome and did not correlate with published topological domains (Figure 5) or syntenic blocks (not shown) near the breakpoint. Two-copy genes that changed expression did fall into clusters of genes related by network interactions.

We projected gene expression changes onto an integrated network model based on expression in published GEO datasets, gene interactions, and protein-protein interactions [31], and observed striking examples of propagating effects in network space. For example, hemizygous *La autoantigen-like (La)* expression was reduced in *Df(2L)ED1315/+* females, as were a number of other genes that are primary (1°) network neighbors of *La* (**Figure 6A**). These genes show enriched expression in ovaries, but are widely expressed in other tissues, and are highly enriched in genes encoding ribosome biosynthetic machinery according to GO term analysis *(p* << 0.001, Holm-Bonferroni corrected Hypergeometric test). This positive relationship between hemizygous expression and expression in network neighbors was more prevalent, but propagation patterns also showed negative interactions. *Suppressor of variegation 205 (Su(var)205)* encodes heterochromatin protein 1 (HP1) that binds H3K9me2/3 and is a general repressor of transcription [32]. Hemizygosity for *Su(var)205* in *Df(2L)ED578/+* females resulted in reduced expression of this negative regulator. The primary neighboring genes connected to *Su(var)205* in the network model showed increased expression (**Figure 6B**), consistent with de-repression when HP1 levels are reduced. We also observed coherent changes in pathways (statistically significant network module clusters [33]) that we could not directly connect to one-copy genes. These were often connected with a few edges to large groups of genes showing the opposite effect. For example, in *Df(2L)ED250/+* females, a strong cluster of genes with ovary-biased expression in wild-type (89% have transcripts enriched in ovary [34]) are down-regulated (**Figure 6C**). These include *Cyclin* genes which are likely expressed in the replicating germline and somatic support cells of the ovary *(CycA, CycB*, and *CycE* [35, 36]), the important female germline transcription factor *ovo* and the known OVO target gene *ovarian tumor (otu)* [37, 38], as well as predicted DSX target genes *Grunge (Gug), domino (dom)*, and the *Insulin receptor (InR)* gene [39]. This cluster of genes is significantly enriched for a host of oogenesis related GO terms and is linked to an even larger cluster of up-regulated genes, 77% of which have oxidative phosphorylation GO terms GO analysis not shown). Energy storage molecules are deposited in the egg by females to support embryonic development and are converted to ATP if not transported to the egg. We suggest that our observations reflect physiological pathway compensation in *Df(2L)ED250/+* females. The effects on two-copy genes with functional rather than physical proximity to one-copy genes strongly suggests that expression changes in the diploid portion of the genome in *Df/+* flies are due to regulatory interactions, not structural changes in the genome or long range contacts.

**Fig 6.**
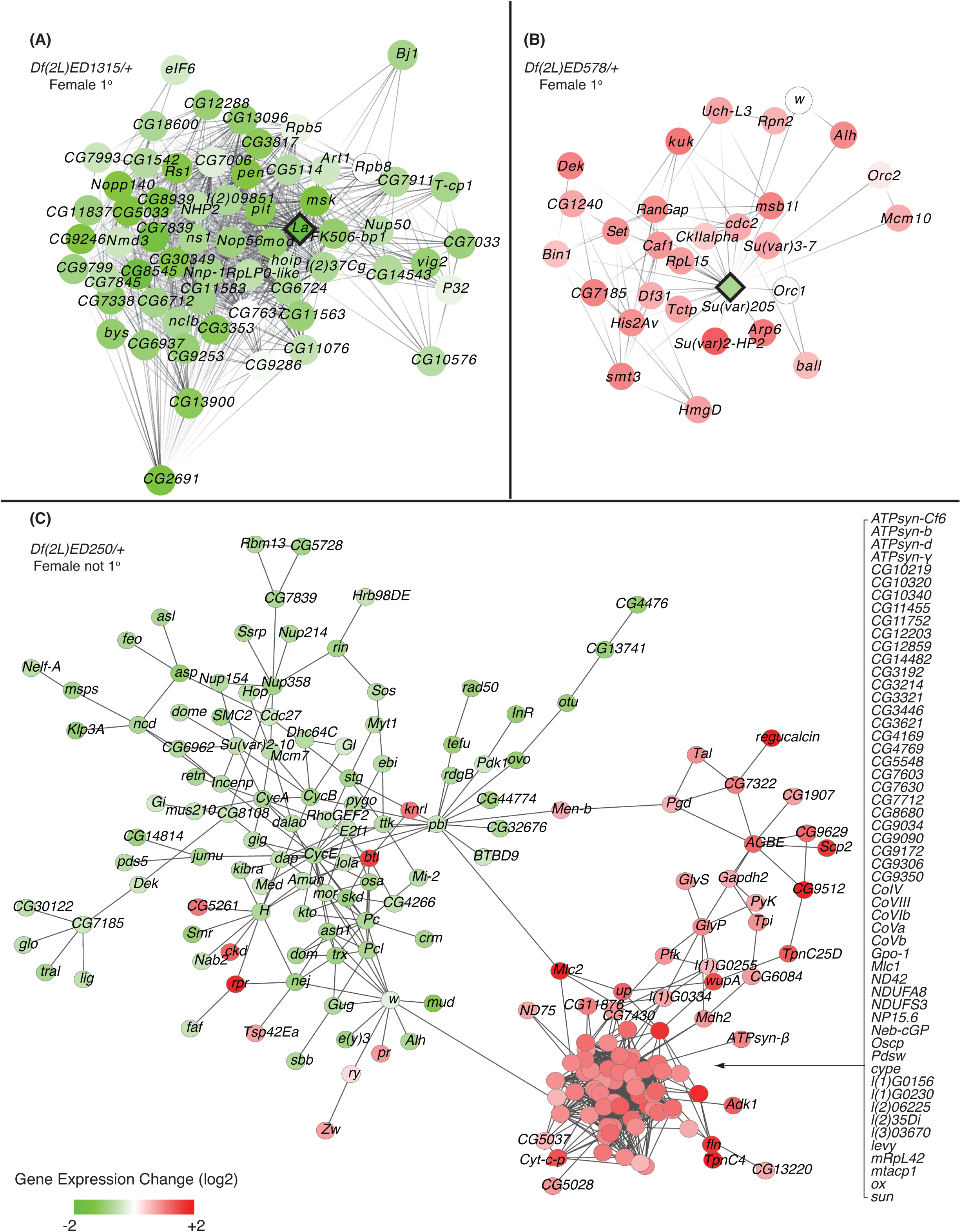
Propagation of gene dose perturbation in gene networks. Nodes represent genes and edges their connections in the integrated network model. A, B) Subnetworks that include *Df* genes (diamond nodes) and their 1° two-copy neighbors (round nodes) in the gene network. Gene expression changes in *Df(2L)ED1315/+* and *Df(2L)ED578/+* females have been projected onto the network. Up-regulated (red) down-regulated (green) gene expression, as well as no change (open), is indicated with shading showing the magnitude of expression change (darker is greater change). C) Identified functional module that is significantly differentially expressed in *Df(2L)ED250/+* females, but with an unknown connection to a one-copy gene(s).

We examined the global relationship between expression of one-copy genes and their two-copy network neighbors in the entire date set (Figure 7) and observed a positive correlation in the expression of one-copy genes and their 1° network neighbors. As network distance increased this correlation degraded and was not distinguishable after 3^°^ connections. This is consistent with a gradual dissipation of the driver perturbation due to the action of corrective feedback responses at each step. We also observed greater expression amplitude in the responding two-copy genes as network distance increased. The one-copy genes and their primary neighbors showed similarly reduced expression, but the 3^°^ neighbors ranged in expression from >50-fold up or down relative to the global control values. These expression relationship patterns were consistent between the sexes and in both the isogenic and hybrid backgrounds.

**Fig 7.**
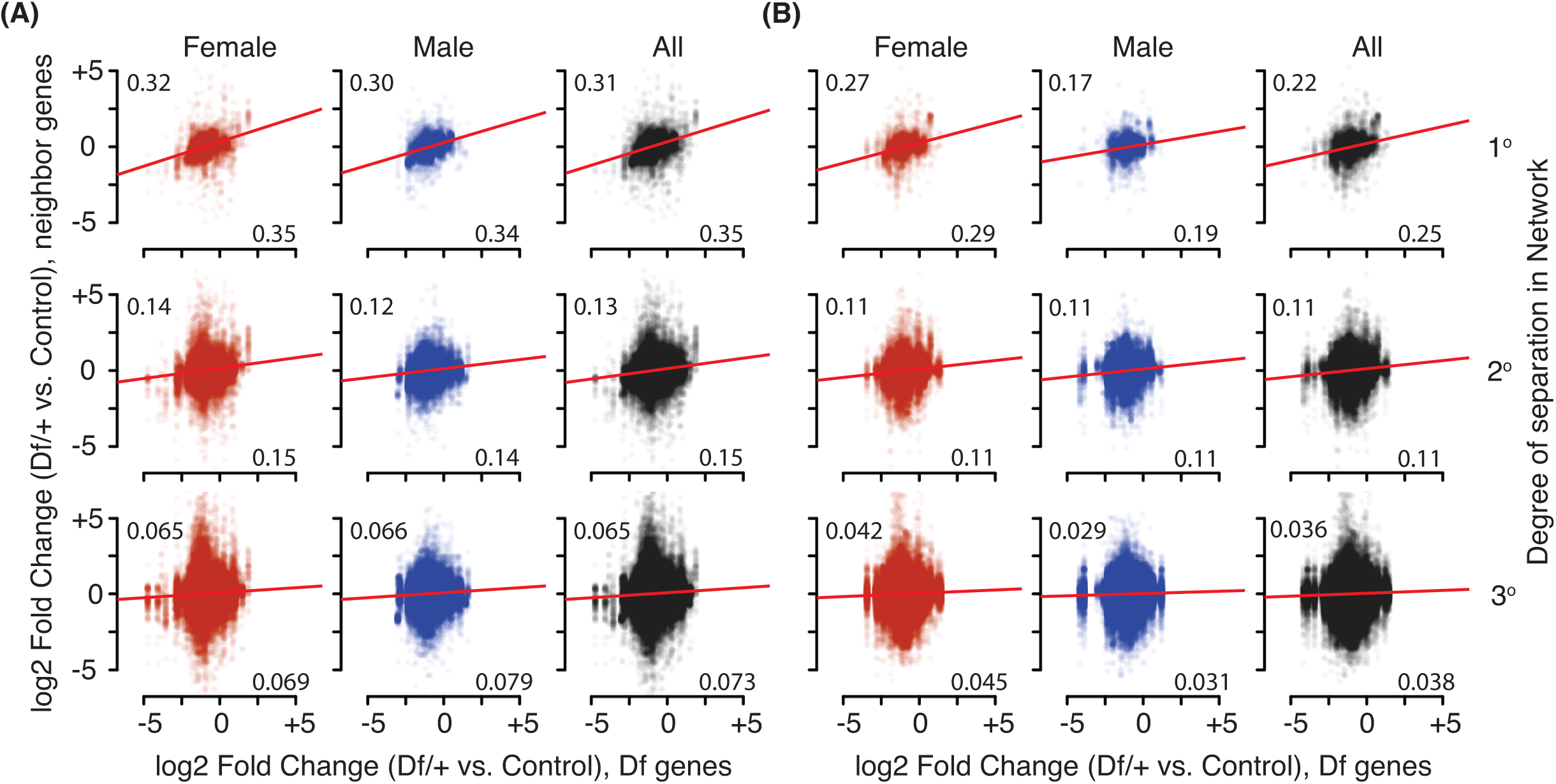
Propagation and dissipation of gene dose perturbation in positively correlating networks. Gene expression changes of two-copy genes that are neighbors of *Df* genes in the integrative network by degree of separation: 1° (primary, top), 2° (secondary, middle) and 3° (tertiary, bottom) are plotted from all *Df* genes (one-copy, x-axis) in females (red), males (blue) and combined (black). Results from both the A) isogenic and B) hybrid background are shown. Red lines are from linear modeling. Pearson’s correlation coefficient (*r*) is shown (upper left in each graph). Bottom-right in each panel displays slopes obtained from linear regression.

We observed similar overall patterns of network propagation and dissipation in both the isogenic and hybrid backgrounds in both sexes. However, the precise genes that changed in response to a given *Df* differed by sex and by genetic background. For example, *Df(2L)ED136/+* males showed many more expression changes than females in both backgrounds (**Figure 8A,B**). In males, other than the one-copy genes, only four genes showed a significant change in both backgrounds. Of the genes showing differential expression in *Df(2L)ED136/+* males, only *CG18600* was also differentially expressed in females. This gene is expressed preferentially in gonads and male accessory gland in wildtype flies [40]. Globally, differently responding genes among the sexes and backgrounds was a strikingly common trend (**Figure 8C,D**). Significant changes in the expression of the one-copy genes showed 27% overlap in the genetic backgrounds, indicating that dose responses were similar among alleles from different backgrounds (also see Figures 2,3). There was a much larger group of two-copy genes that showed significant changes in gene expression. In our analysis of chromosome *2L* we observed that 10,418 genes display significant changes at least once in any of the test genotypes (76% of genes). In striking contrast to similarity in dose responses among one-copy genes, the genes responding to the perturbation were usually different. We observed < 1% overlap between backgrounds *(p* = 0.39 to 0.61) and even fewer genes showed changed expression in both sexes and in both backgrounds. This is perhaps unsurprising given that there are ~600K heterozygous SNPs and Indels in the hybrid background relative to the isogenic background (**Supplemental file 1**) and given the pervasive sex-bias in *Drosophila* gene expression. Thus, while there are coherent pathway responses to hemizygous driver perturbations, the exact path through network space was highly dependent on sex and genetic background.

**Fig 8.**
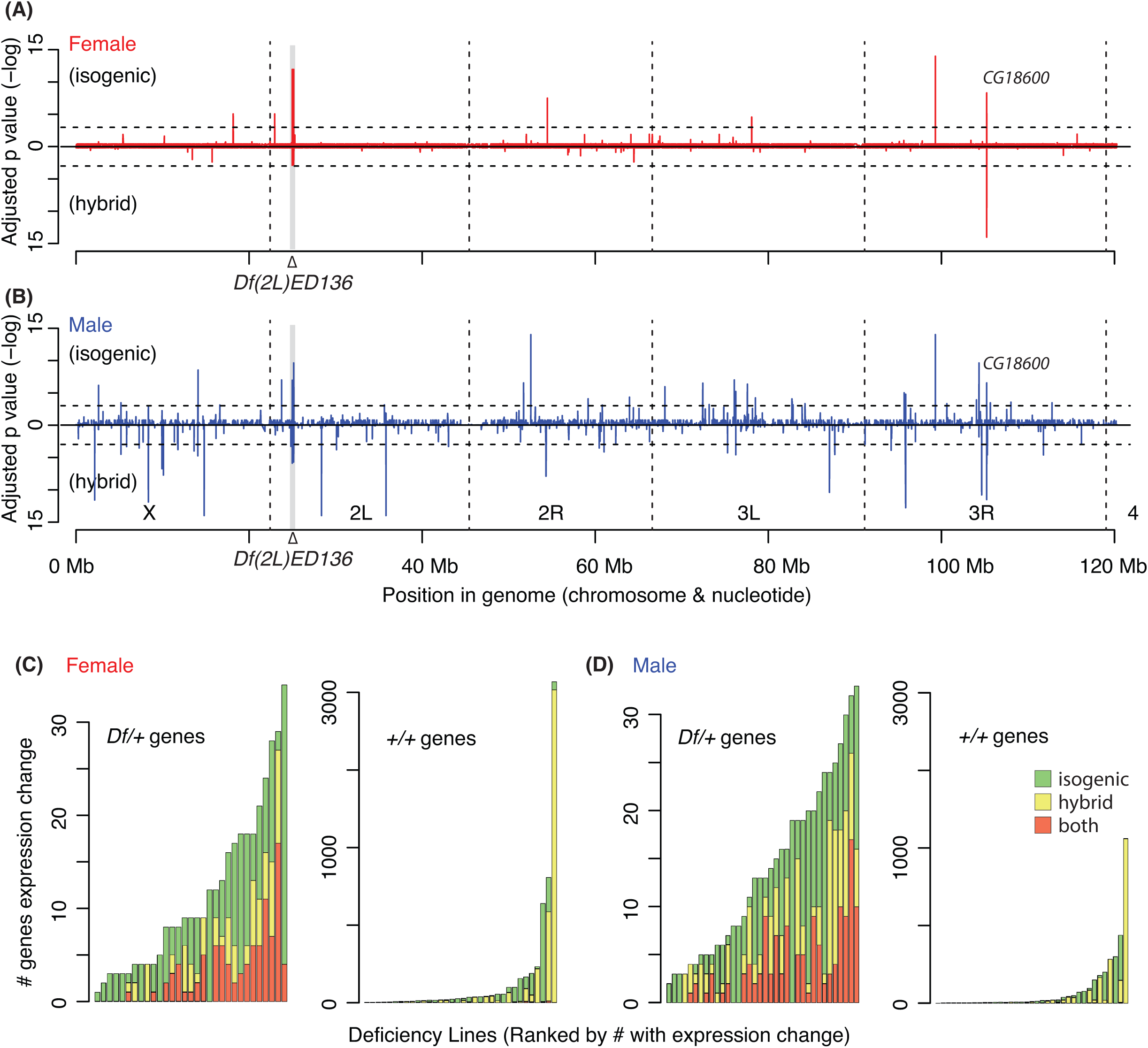
*Df/+* drivers cause genome-wide gene expression change in a genetic background-dependent manner. A, B) Adjusted *p* values of gene expression change (-log scale) across the genome for *Df(2L)ED136/+* females (A) and males (B). *Df* regions (grey) *p* value = 0.05 (horizontal dotted lines), chromosome arms (vertical dotted lines), and genomic position (scale) are shown. C, D) Stacked plots of significantly differentially expressed one copy (left) and two copy (right) genes. Note the two scales of numbers of differentially expressed genes. Numbers of genes in the isogenic (green), hybrid (yellow), or both genetic backgrounds (red) are shown.

## DISCUSSION

Large deletion collections are very widely used tools by the *Drosophila* research community, representing the most commonly ordered stocks from the Bloomington Drosophila Stock Center. Unfortunately, we know very little about the effects these deletions have on the global fly transcriptome and here we describe our initial efforts to address this issue. We have touched on three aspects of the effects deletions have on the transcriptome: 1) the primary effect of hemizygousity on gene expression; 2) juxtaposition of regions of the genome that are normally distant; and 3) the effects of deletions at a distance in either network or physical space.

### Dosage effects and compensation

In a deficiency, there is expression from only one allele of the genes within the aneuploid deleted region, and while this generally results in expected reductions in gene expression, the response of an individual gene to copy number reduction is highly dependent on feedback regulation [8] and buffering, both of which are inherent properties of biochemical pathways [8, 10, 12, 13].

Different degrees of autosomal compensation have been reported for *Drosophila melanogaster*, ranging from no compensation to nearly 2-fold up-regulation of hemizygous gene expression [9–11, 13–15, 41]. Some of the differences in compensation values are probably due to biology, such as the varied responses of aneuploid tissue culture cells [15]. However, data compression in microarray-based studies also contributes to over estimating dosage compensation especially at low expression levels where array responses are nonlinear [42–44]. For example, a microarray study where stringent expression cutoffs were applied to measure compensation levels [10] resulted in the same 1.1-fold compensation from the hemizygotic genes we report here. Reanalysis with the same stringent method (not shown) results in 1.1-fold compensation rather than the 1.4-fold compensation reported in another microarray study [8]. Our low expression cutoff in this study does not underestimate compensation, as the extensive set of spike-in RNAs we analyzed shows little evidence of data compression (see Methods).

There has been debate about whether there is a regional response to reduced gene dose in *Drosophila* [8, 10, 12, 13]. In our analysis of chromosome *2L*, we found that many genes showed poor or partial compensation, while others showed excellent dosage compensation, which is consistent with feedback and buffering models. Given the propagation of gene expression changes from one copy segments to two copy genes through regulatory network connections, it is clear that gene dosage perturbs gene networks. Thus, we suggest that traditional gene regulation involving feedback can explain the vast majority of the dosage compensation response on chromosome *2L*. However, we also identified blocks of well compensated or over compensated genes in females and could correlate these with repressive chromatin domains. Such chromatin-based responses to autosomal aneuploidy are analogous to the chromosome-wide MSL and POF systems that alter chromosome wide expression from the *X* and the ancestral *X* (the current chromosome 4) in *Drosophila* [45].

Interestingly, the chromatin domains resulting in superior compensation were repressive, with diagnostic LamB and/or PcG enrichment. The PcG proteins can mediate pairing-dependent silencing [46], which has the counter-intuitive effect of increasing expression of one-copy genes. We observed the same effect in some clusters of one-copy genes in this study. In *C. elegans*, the two *X* chromosomes in *XX* hermaphrodites are down-regulated to counteract the increased *X* chromosome expression that *X0* males use to equilibrate *X* and autosomal gene expression [47]. *XX* down-regulation is achieved by strengthening the attachment of both *X* chromosomes to the repressive regions while this is relaxed in males with one-copy of the *X* to increase expression [48–50]. The increased repression of two-copy genes relative to one-copy genes due to deletions in LaminB domains is similar, suggesting a plausible model for the evolution of *X* chromosome dosage compensation. Genetic material, including pairing-dependent repressive domains are progressively lost from neo*-Y* chromosomes as they diverge from the *X* homolog on evolutionary timescales. The loss of pairing dependent repressive domains could lead to regional dosage compensation prior to evolution of a specific *X* chromosome-wide mechanism.

Curiously, we observed this regional LaminB and PcG dosage compensation response only in females. On the one hand this may reflect ascertainment bias since we used domains defined by work in tissue culture cells and embryos [24–26] and it is possible that the arrangement of domains in the adult fly could be significantly different and sex-biased. On the other hand, sex-specific differences in the nature of heterochromatin have also been noted [51] and it is possible that there is a general weakening of repressive domains in males, reducing the possibility for regional autosomal dosage compensation due to further de-repression. The fact that a group of well-compensated genes was only found in females, regardless of where they were located, favors a female-biased derepression model. Clearly, additional experiments will be required to investigate this curious finding.

### Breakpoints

*Df* breakpoints bring two regions of the genome together that are usually distant in the linear chromosome. This can result in breakpoint proximal changes due to transcription unit fusion as occurs in many cancers and has been especially well studied in immune cell tumors [52]. However, we observed only one such case in our analysis: *Df(2L)ED680* results in a fusion transcript of *taiman* and mini*-white* (the marker for deletion). It is also clear that some genes have enhancers and silencers located many kb from the promoter [53–55]. It is also clear that the genome is organized into chromatin domains flanked by insulator regions, which could facilitate regional transcriptional control [56]. Deficiencies delete insulator sites resulting in the creation of novel arrangements of insulator pairs. If this creates a novel gene expression regulatory milieu, then transcription should be altered. Our analysis indicates that across ~ 20 Mb of the genome we surveyed, the vast majority of the regulatory information is within the gene body or ~100 bp upstream. This agrees with work where inversions generated within *Drosophila* neighborhoods of co-expressed genes failed to disrupt co-regulated gene expression [20]. It is possible that there are highly deleterious cases where generation of a *Df* is dominant lethal, but even this is likely to be rare. In a study that generated a large number of FRT (Flippase Recognition Target) deletions, 6% of the pairs failed to produce a deletion [57]. The majority of regions can be joined without dominant lethality. We suggest that effects of DNA topological domains and long-range enhancer promoter interactions are rare in *Drosophila* adults.

### Network interactions

We observed substantial changes in gene expression throughout the genome, not just in the one copy regions, and these are likely due to “error” propagation as is expected in a dynamic biological system. The primary two copy network neighbors of one copy genes change expression in response to reduced dosage of genes uncovered by deletions. In many cases *Dfs* reveal tightly connected subnetworks of genes expressed in a particular tissue, such as the gonads, gut, or eyes. This is unsurprising, since dose-dependent enhancers and suppressors have been identified by screening with deletions for decades. Such dominant genetic interactions are very valuable for finding near neighbors in genetic pathways. For example, much of what we learned about the germline sex determination pathway in *Drosophila* began with screens identifying pairs of interacting genes: *sans fille* (*snf*) and *fl(2)d* loci show dominant interactions with *Sex-lethal (Sxl)* resulting in germline tumors [58, 59], and are now known to be components of the splicing machinery participating in *Sxl* autoregulation [60]. Additionally, screening *Df* heterozygotes for dominant interactions with *ovo^D^* identified a number of interacting genes [61], including the direct OVO target, *otu* [38]. Interestingly, we find that particular *Dfs* result in coordinate changes in *ovo* and *otu* expression, raising the possibility that one could screen the *Df* kit directly and then predict the genetic interactions.

## MATERIALS and METHODS

### Fly lines

*Drosophila melanogaster* were raised at 25°C on the standard yeast/cornmeal medium (Fly Facility, University of Cambridge, UK) for *w^1118^* isogenic flies, and on the standard fly agar (Bloomington Drosophila Stock Center, Indiana University, IN) for *w^1118^/OregonR* hybrid flies. Virgin *w^1118^* or ModENCODE *OregonR* females were crossed to DrosDel strain males and the non-balancer progeny were collected and aged for 3–5 days. Progeny were allowed to mate freely.

We used the following DrosDel lines: *Df(2L)ED105, Df(2L)ED1050, Df(2L)ED1054, Df(2L)ED108, Df(2L)ED1109, Df(2L)ED1161, Df(2L)ED1196, Df(2L)ED1200, Df(2L)ED1202, Df(2L)ED1203, Df(2L)ED1226, Df(2L)ED123, Df(2L)ED125, Df(2L)ED12527, Df(2L)ED1272, Df(2L)ED1303, Df(2L)ED1305, Df(2L)ED1315, Df(2L)ED1317, Df(2L)ED134, Df(2L)ED136, Df(2L)ED1378, Df(2L)ED1454, Df(2L)ED1466, Df(2L)ED1473, Df(2L)ED19, Df(2L)ED206, Df(2L)ED334, Df(2L)ED343, Df(2L)ED347, Df(2L)ED353, Df(2L)ED354, Df(2L)ED369, Df(2L)ED384, Df(2L)ED385, Df(2L)ED40, Df(2L)ED441, Df(2L)ED4559, Df(2L)ED4651, Df(2L)ED475, Df(2L)ED479, Df(2L)ED489, Df(2L)ED49, Df(2L)ED50001, Df(2L)ED690, Df(2L)ED695, Df(2L)ED700, Df(2L)ED7007, Df(2L)ED729, Df(2L)ED746, Df(2L)ED761, Df(2L)ED773, Df(2L)ED774, Df(2L)ED775, Df(2L)ED776, Df(2L)ED7762, Df(2L)ED777, Df(2L)ED778, Df(2L)ED779, Df(2L)ED784, Df(2L)ED7853, Df(2L)ED793, Df(2L)ED80, Df(2L)ED800, Df(2L)ED8142, Df(2L)ED8185, Df(2L)ED87, Df(2L)ED929*, and *Df(2L)ED94* were analyzed for both the hybrid *(w^1118^/OregonR)* and the isogenic (*w^1118^*) backgrounds. *Df(2L)ED1102, Df(2L)ED1231, Df(2L)ED21, Df(2L)ED243, Df(2L)ED247, Df(2L)ED250, Df(2L)ED270, Df(2L)ED279, Df(2L)ED280, Df(2L)ED284, Df(2L)ED285, Df(2L)ED292, Df(2L)ED299, Df(2L)ED3, Df(2L)ED320, Df(2L)ED33, Df(2L)ED499, Df(2L)ED501, Df(2L)ED508, Df(2L)ED548, Df(2L)ED578, Df(2L)ED5878, Df(2L)ED611, Df(2L)ED62, Df(2L)ED623, Df(2L)ED629, Df(2L)ED647, Df(2L)ED6569, Df(2L)ED678*, and *Df(2L)ED680* were analyzed only in the *w^1118^* isogenic background, and *Df(2L)ED1004, Df(2L)ED632*, and *Df(2L)ED8186* were analyzed only in the *w^1118^/OregonR* hybrid background. *Df(2L)ED1050, Df(2L)ED123*, and *Df(2L)ED611* showed no clear reduction in hemizygous gene expression raising the possibility that they are not deletions, although they were homozygous lethal. Additionally, *Df(2L)ED1050* and *Df(2L)ED123* complemented mutations that they should uncover, suggesting that they are not correctly identified *Dfs*. Inclusion/exclusion of these three *Dfs* did not alter overall compensation values at the rounding levels reported here. We excluded them from one-copy analysis, but they serve as additional controls.

### RNA-Seq molecular biology

Preparation of RNA sequencing libraries for the *w^1118^* isogenic background is described in Cho *et al*. (Companion Paper). For the hybrid background files, single, day 3–5 adult male or female flies were partially crushed, stored in 100ul of RNAlater (Life Technologies, Grand Island, NY), and frozen at -80°C for long-term storage until RNA preparation. RNA was isolated from each genotype in biological triplicates.

We used Mini-BeadBeater 96 (Biospec Products, Bartlesville, OK) for the homogenization of flies. Approximately 100μl of 1mm glass beads (Biospec Products, Bartlesville, OK) were added to the flies in RNAlater in 1ml Axygen 96 well plate (Corning, Union City, CA). We processed plates 3 x 1min with a 2min rest on ice between each homogenization. We added 600μl of RLT buffer (Qiagen, Valencia, CA) to each well to dilute the RNAlater solution. Total RNA was isolated with RNeasy 96 kits (Qiagen, Valencia, CA) according to manufacturer’s handbook (Protocol for Isolation of Total RNA from Animal Cells using spin technology, Cat#19504). The amount of total RNA extract was measured using Quant-iT RiboGreen (Life Technologies, Grand Island, NY).

We mixed 400ng of total RNA in 50μl of nuclease-free water with 50μl of 2:5 dilution of Dynabeads Oligo(dT)25 (Life Technologies, Grand Island, NY) that we pre-rinsed and diluted with Binding Buffer (20mM Tris-HCl pH7.5, 1.0M LiCl, 2mM EDTA). We heated the mixture to 65°C for 5min in a thermocycler, and cooled down on ice for 1min. After 15min of incubation at room temperature, we collected the beads with a magnetic stand, and rinsed with 200μl Washing Buffer (10mM Tris-HCl pH7.5, 0.15M LiCl, 1mM EDTA) for 1min at 1,000 rpm (Thermomixer, Eppendorf, Hauppauge, NY). We collected the beads again with a magnetic stand, and eluted with 50μl of 10mM Tris-HCl pH7.5 at 80°C for 2min. We rebound the eluate to the beads by incubating with 50μl Binding buffer, and rinsed with 200μl Washing Buffer as above. We eluted and fragmented the poly A+ RNA with 16μl of Fragmentation Buffer that contained 1:4 dilution of 5X First Strand Buffer from Protoscript II (New England BioLabs, Ipswich, MA), 500ng of random primers (Life Technologies, Grand Island, NY), and 20pg of ERCC spike-ins Pool 78A or 78B [22, 62] at 95°C for 6min. The beads were removed using a magnetic stand, we reverse transcribed with10units of SuperRase-in (Life Technologies, Grand Island, NY), 100units of Protoscript II reverse transcriptase, 5mM DTT, and 625 μM dNTPs (Enzymatics, Beverly, MA). We used a thermocycler set at 25°C for 10min; 42°C for 50min; 70°C for 15min. We cleaned-up the DNA-RNA hybrids with 1.9 volumes of MagNA beads[63], and mixed with 0.85 volumes of ethanol and we bound DNA-RNA hybrid to the beads at room temperature for 15min. We collected beads on a magnetic stand and rinsed with 200μl of 80% ethanol twice. After air-drying for 5min, we eluted the beads with 16μl of Elution Buffer. For the second strand synthesis, we added 2.5 units of RNase H (Enzymatics, Beverly, MA.) and 10 units of DNA polymerase I in 1X Blue Buffer (Enzymatics, Beverly, MA.), 10mM DTT, 0.5mM each dATP, dCTP, and dGTP (Enzymatics, Beverly, MA.) and 1mM dUTP (ThermoFisher Scientific, Waltham, MA. 1mM final) and incubated for 5 hours at 16°C. We cleaned-up DNA products MagNA beads and eluted as above but we used bound beads for the next purification steps. Purified double-stranded DNA was subjected to end-repair with NEBNext End Repair Module (New England BioLabs) as per manufacturer’s protocol for 20μl samples. We replenished beads with 1.9 volume of XP buffer [64] and cleaned up as above. For adapter ligation, we performed adenylation on blunt-ended DNA by adding 2.5 units of Klenow 3’-5’ exo (Enzymatics, Beverly, MA) and incubation at 37°C for 30min in 1X Blue Buffer. We again used 1.9 volume of XP buffer to clean up as above. We eluted DNA with 10μl of Elution buffer, and added 1μl of one of 24 differently barcoded adapters from the TruSeq v2 kit (Illumina, San Diego, CA) for 24μl ligation reactions with T4 DNA ligase (Rapid) (Enzymatics, Beverly, MA.) for 20min at 20°C. We stopped the ligation by adding volume of 0.03M EDTA, and cleaned up with 24μl of XP buffer as above. After eluting with 30μl Elution buffer, we cleaned up again with 1 volume of XP buffer add brought samples to a 12μl final volume. We digested dUTP incorporated strands of DNA with 5 units of Uracil DNA Glycosylase (New England BioLabs) for 30min at 37°C. Then we amplified with 1.5μl of P5 and P7 primers (Integrated DNA Technology, Coralville, Iowa) with KAPA HiFi HotStart DNA polymerase (KAPA biosystems, Wilmington, MA) in 30ul reactions. We used the following PCR parameters: 98°C for 45 sec, 14 X 98°C for 15sec; 60°C for 30sec; 72°C for 30 sec, followed by 72°C for 5min. We cleaned up amplified DNA with MagNA beads. Libraries were quantified with Quant-iT PicoGreen (Life Technologies), and pooled to be sequenced in Illumina HiSeq 2000 (Illumina, San Diego, CA).

### RNA-Seq data analysis

RNA-Seq results were analyzed as in Cho et al. (**companion paper**) with a minor difference in handling the ERCC spike-ins. Briefly, the short reads generated from the analysis were mapped onto the *Drosophila* reference genome (Release 5, with no “chrU” and “chrUextra” scaffolds) using TopHat 2.0.11 [65]. We used Cufflinks 2.2.1 [66] to measure gene expression levels in Fragments per Kilobase per Million mapped reads (FPKM). We also measured FPKM values from intergenic regions as in [14, 67], which we used to determine expression cutoff levels as 0.6829118 for the isogenic background results, and 0.8140542 for the hybrid genetic background (see below). We used HTseq 0.6.1p1 [67] [68] and “voom” in the R limma package [68] to obtain raw reads counts at the gene level, and to call differential gene expression as in (Cho et al, companion paper). Benjamini-Hochberg multiple hypothesis correction of the P values was used throughout the manuscript. FPKM values for the spike-ins were separately determined as abundant transcripts influence of estimation of gene expression. In the calculation, the number of reads from both genes and spike-ins, not solely from spike-ins, was used as the denominator of FPKM to infer the lowest detectable expression levels of genes. In describing gene expression changes, we report expression values for genes that produce polyA+ mRNA in the gene model, since we followed a poly-A purification protocol. For example, we detected expression of histone transcripts, but the values were highly variable between libraries due to their lack of poly-A tails, and essentially followed the values of residual rRNAs in the sequencing libraries [69]. Expression results presented were robust to normalization methods used in different bioinformatics tools (FPKM, DESeq, and TMM). We used Expectation-Maximization method to identification of the fully compensated group of genes in Figure 4 using “mclust” package in R (**doi:10.1007/s00357-003–0015–3**). In the analysis, two ellipsoidal distributions of equal orientation, “EEE”, were used as models for the clustering algorithm.

We used TopHat 2.0.11 to identify fusion transcripts. Potential fusion transcripts that have at least 15 bp long anchor sequences were surveyed. For intra-chromosomal fusion events, we investigated potential fusion transcripts with more than 683 bp distances between juxtaposed regions based on the shortest length of *Dfs* (fusion-anchor-length 15 and fusion-min-dist 683 parameters).

We observed outstanding biological replicate profiles (Pearson’s r <0.9) in all 396 isogenic samples (**Figure 9A**). However, we observed 10% outliers in the single fly profiles, and therefore we removed all samples where Pearson’s r <0.9, and we used the two duplicates with the highest correlation. The sexual identify of a sample is self-reported in the expression profile (**Figure 9B,C**). Detection of low-level gene expression is complicated by the contributions of noise, which vary between libraries. To mitigate this problem we measured reads from intergenic regions (trimmed to account for variation in transcription start and stop sites) and determined the 95th percentile as a low expression cutoff (**Figure 9D**). While some of this intergenic expression may be due to strain-specific transcripts or un-annotated genes, much is likely to be due to noise such as ectopic Pol-II initiation, inclusion of contaminating genomic DNA, sequencing, and/or mapping errors. Ratios and data compression measurements are critical for dosage compensation analysis. We used pools of ERCC controls in each sample library to produce 1.5:1, 1:1, and 1:1.5 ratios across a > 2^15^ input concentration range (**Figure 9E**). Ratio measurements show a clear increase in scatter when input was low. However, even at very low input, there was only modest compression, and no evidence of compression in the useful range for this work.

**Fig 9.**
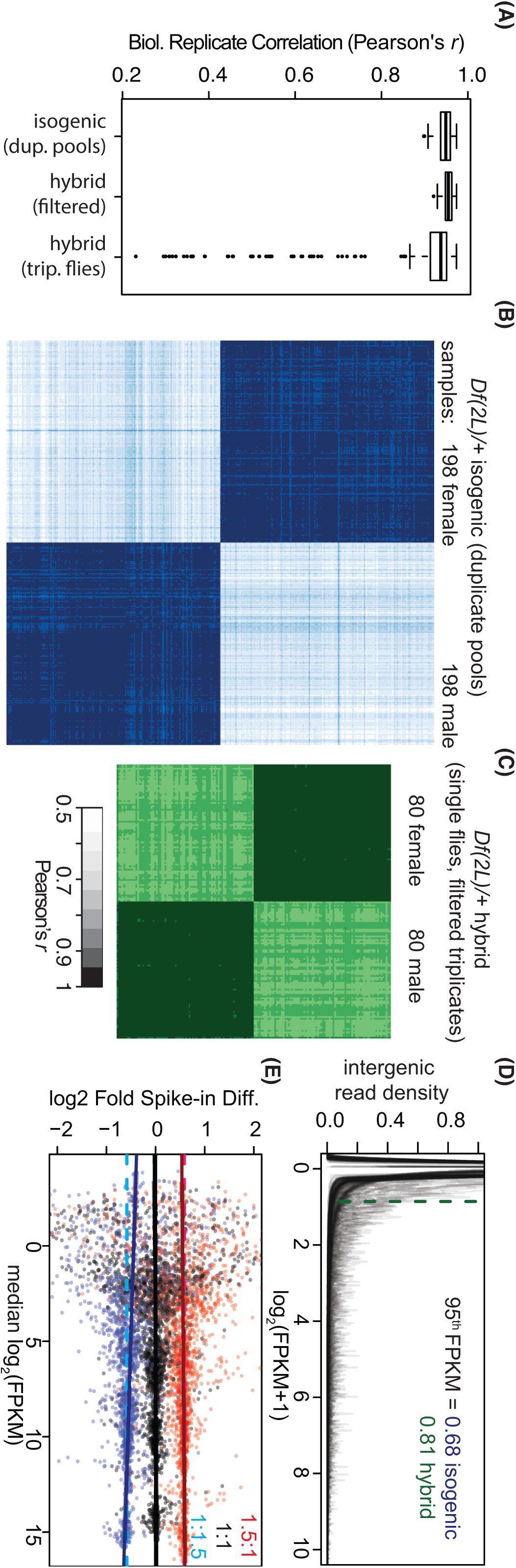
Data quality analysis. A) Boxplots of Pearson’s correlation coefficient (*r*) between experimental replicates. For replicates from the hybrid genetic background we applied a filter at *r* < 0.9 and selected best duplicates from triplicated experiments to remove outliers. B, C) All sample-to-all sample pairwise comparisons (darker is higher Pearson’s *r*). D) RNA-Seq signals from intergenic regions collectively overlaid as density plots across signal in FPKM. Median of the top 95 percentile in the distribution is indicated (dotted lines). E) Signals from ERCC spike-in sets having 1.5:1 (red), 1:1 (black), and 1:1.5 (blue) ratios between libraries as indicated. Expected ratios (solid lines) and trend lines from linear modeling (dotted) are shown.

### Detection of sequence variants between *OregonR* and *w^1118^* flies

DNA-Seq reads of the Bloomington Stock Center *w^1118^* line was obtained from the Sequence Read Archive (SRR630490). We mapped the raw reads to the reference genome using Bowtie 2 [70] with default parameters. We used Samtools to call SNPs from the mapping result [71]. The calls were filtered to have equal to or more than quality score 20 (-Q 20). Any calls that have more than twice the average read depth were discarded. Based on the mapping, we incorporated substituted bases, or SNPs, as previously described [72] to have “*w^1118^* SNP-substituted genome”. The differences between *w^1118^* SNP-incorporated genome and *OregonR* DNA-Seq results were identified using Samtools as described above.

## Data Access

The gene expression profiles generated in this study are available in GEO with accession numbers of GSE61509 (isogenic genetic background) and GSE73920 (hybrid genetic background).

## Author Contributions

CW, RE, MP, JR, TK, KC, SR, and BO performed Drosophila crosses and collected flies. KC performed complementation tests of *Dfs*. HL, CW, SR and BO prepared samples. HL and CW prepared sequencing libraries and processed sequence data. HL, DYC, TMP, and BO performed analysis. HL, DYC, SR, TMP and BO prepared the manuscript. All authors provided feedback, and read and approved the final manuscript.

## Acknowledgements

This research was supported in part by the Intramural Research Program of the NIH, The National Institute of Diabetes and Digestive Diseases (BO) and the National Library of Medicine (TMP). HL was supported by Korean Visiting Scientist Training Award (KVSTA, HI13C1282). Our research utilized the high-performance computational capabilities of the Biowulf system at the NIH, Bethesda, MD. Stocks obtained from the Bloomington Drosophila Stock Center (NIH P40OD018537) were used in this study. Certain commercial equipment, instruments, or materials are identified in this document. Such identification does not imply recommendation or endorsement by NIH.

